# Whole genome sequencing of a wild yam species *Dioscorea tokoro* reveals a genomic region associated with sex

**DOI:** 10.1101/2022.06.11.495741

**Authors:** Satoshi Natsume, Hiroki Yaegashi, Yu Sugihara, Akira Abe, Motoki Shimizu, Kaori Oikawa, Benjamen White, Aoi Kudoh, Ryohei Terauchi

**Author notes:** Correspondence to Ryohei Terauchi.

## Abstract

*Dioscorea tokoro* is a wild species distributed in East Asia including Japan. Typical of the genus *Dioscorea, D. tokoro* is dioecious with male and female flowers borne on separate individuals. To understand its sex determination system and to serve as a model species for population genomics of obligate outcrossing wild species, we set out to determine the whole genome sequence of the species. Here we show 443 Mb genome sequence of *D. tokoro* distributed over 2,931 contigs that were anchored on 10 linkage groups. Linkage analysis of sex in a segregating F1 family revealed a sex determination locus residing on Pseudochromosome 3 with XY-type male heterogametic sex determination system.

## Introduction

The genus *Dioscorea* belongs to the monocotyledons and has 450 - 600 species distributed mainly in tropical and subtropical area of the world (Coursey, 1972; Sugihara et al. 2021). Cultivated species of *Dioscorea* are collectively called yam, which includes Guinea yam (*D. rotundata*) of West Africa accounting for more than 90% of the world yam production (FAOSTAT, 2018). The entire genus of *Dioscorea* is characterized by dioecy, with male and female flowers borne on separate individuals. Consequently, the species of *Dioscorea* have obligate outcrossing, resulting in a higher level of heterozygosity and frequent inter-species hybridization (Terauchi et al. 1992; Chaïr et al. 2010, 2016; Girma et al. 2014; Scarcelli et al. 2006, 2017; Siadjeu et al. 2018; Sugihara et al. 2020, 2021; Bredeson et al. 2022). Previously we reported the whole genome sequence of *D. rotundata* with 570 Mb in size (Tamiru et al. 2017), which served as a reference to study population genomics of *D. rotundata* and its wild relatives to reveal the origin of Guinea yam (Scarcelli et al. 2019; Sugihara et al. 2020).

*Dioscorea tokoro* is a wild species belonging to the section Stenophora. It is widely distributed in East Asia including Japan. *D. tokor*o is a diploid species with a chromosome number 2n = 2x = 20. It is a perennial species with rhizomes. In spring, shoots emerge from rhizomes and develop to vines that twine around nearby trees in an anti-clockwise direction which expand alternate leaves (**Fig 1**). The species commonly occurs along the fringes of forests in Japan. Crossing experiment is easy and the generation time is relatively short (1-2 years), so that the species has been serving as a model species of *Dioscorea*. The species was subjected to studies of population genetics (Terauchi 1990:Terauchi and Konuma 1994; Terauchi et al.1997), linkage mapping and elucidation of sex determination mechanisms (Terauchi and Kahl, 2004).

**Fig. 1.**
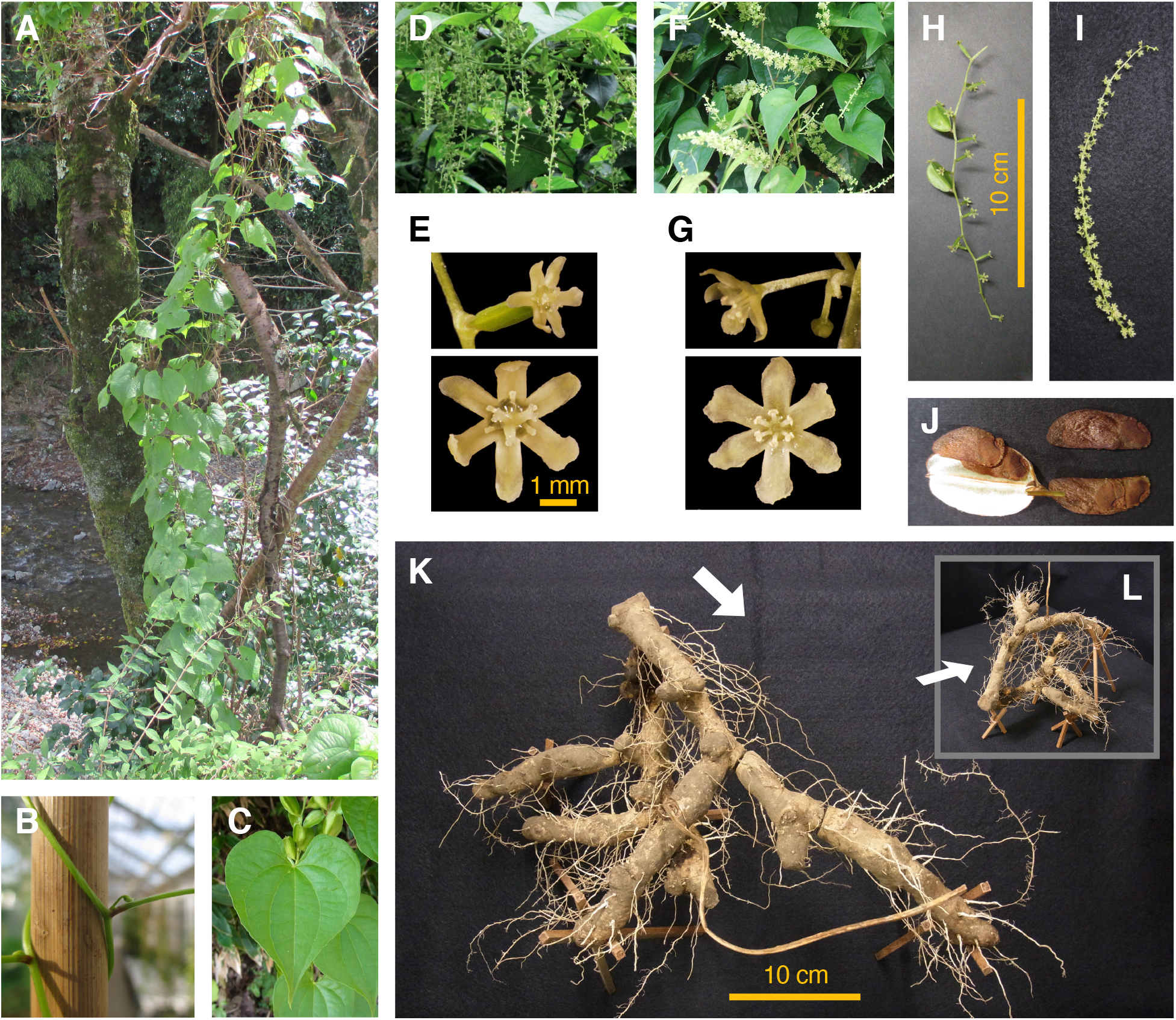
Botanical characteristics of *Dioscorea tokoro*. (A) *D. tokoro* is a herbaceous climber species. Aerial stems twine around tree trunks. (B) Stem twines in an anti-clockwise direction (left-handed; sinistrorse). Leaves alternate. (C) Leaf shape is usually heart-shaped. Leaf blades are typically 5-12 cm long and 5-12 cm wide. (D) Female pendulous inflorescences. (E) Close-up view of a female flower. Three-locular ovary are below the petal. Three-lobed pistil and six degenerated stamens around the pistil are seen. Petal apex is round and curled inward. (F) Male upright inflorescences. (G) Close-up view of a male flower. Pedicel branches from the base and has a few flowers. Six stamens, and degenerated pistil in the center. Petal apex is round and curled inward. The scale is same as (E). (H) Female inflorescence with immature obovate-elliptic capsules. Capsules reflex and dehisce at maturity. (I) Male inflorescence. The scale is same as (H). (J) Mature fruit has three capsules, with winged two seeds placed alternately overlapped near its base. Seed’s wing is biased wider toward capsule apex. (K) Underground rhizome of *D. tokoto*. (L) A side view of the rhizome from a different angle. The direction of the white arrow corresponds to (K).

To serve as a platform for future genomics study of the species, we here report the whole genome sequence of *D. tokoro*. We combined Oxford Nanopore long read sequences and Illumina short read sequences for de novo assembly to generate contigs. The contigs were further anchored on linkage maps to generate pseudochromosomes. RNA-seq data were used for gene prediction. A putative locus involved in *D. tokoro* sex determination was identified on Pseudochromosome 3.

## Results

### Estimation of size of *D. tokoro* genome by flow cytometry

We used a *D. tokoro* individual Kita1 collected at Kitakami, Iwate, Japan as well as *D. rotundata* accession TDR96-F1 with known genome size (∼570 Mb, Tamiru et al. 2017), as the material for flow cytometry (FCM) analysis using nuclei prepared from fresh leaf samples. DNA of isolated nuclei were stained with propidium iodide and analyzed by a flow cytometer. The value of G1 peak mean of *D. tokoro* was 206.5, whereas that of *D. rotundata* was 303.6. The ratio between the two species was 0.68 (206.5/303.6). From these values the genome size of *D. tokoro* was estimated to be ∼388 Mb (570 Mb × 0.68) (**Fig. S1**).

### Reference assembly using Oxford Nanopore Technology

Genomic DNA was extracted from fresh leaves of *D. tokoro* Kita1 and subjected to Oxford Nanopore Technologies (ONT) sequencing. As a result, we obtained a total of 2,515,235 reads amounting 27.4 Gb in size (**Table S1**). We also performed Illumina sequencing of 35 - 251 bp read-length (total 24.6 Gb; obtained by MiSeq) as well as 150 bp read-length (total 37.8 Gb; obtained by HiSeq4000) (**Table S2**). We assembled ONT reads and Illumina sequence reads using a hybrid assembler MaSuRCA v3.3.4 (Zimin et al. 2013) with Flye assembler v2.6 (Kolmogorov et al. 2019) running internally, which generated *D. tokoro* draft genome sequence consisting of 2,931 contigs amounting 443.5 Mb with N_50_ being 586,368 bp (**Table 1**). The estimated genome size by *k*-mer analysis of the reads was 438.7 Mb. These estimated genome size based on DNA sequencing were larger than 388 Mb as estimated by the FCM analysis.

**Table 1.** Summary of a reference genome of *D. tokoro* (Kita1).

|  |  |
| --- | --- |
| Total number of contigs | 2,931 |
| Total base-pairs (bp) | 443,501,147 |
| Estimated genome size from k-mer (bp) | 438,704,233 |
| Average contig size (bp) | 151,313 |
| Longest contig (bp) | 6,172,819 |
| Shortest contig (bp) | 502 |
| N50 (bp) | 586,368 |
| Total BUSCO groups searched | 1,614 |
| Complete BUSCOs (%) | 98.0 |
| Complete and single-copy BUSCOs (%) | 92.9 |
| Complete and duplicated BUSCOs (%) | 5.1 |
| Fragmented BUSCOs (%) | 1.1 |
| Missing BUSCOs (%) | 0.9 |

### Anchoring of contigs on *D. tokoro* linkage maps

To generate *D. tokoto* pseudochromosomes, we mapped the contigs on ten linkage groups. For this purpose, we crossed a female individual Waka1 (P1) with a male individual Kita1 (P2) to obtain F1 progeny comprising 186 individuals (**Fig. S2**; **Table S3**). These plants were genotyped by RAD markers (**Fig. S3**; Baird et al, 2008). We identified 946 SNPs and 180 presence/absence polymorphisms (PAs) that are heterozygous in P1 and homozygous in P2 parents, and 724 SNPs and 880 PAs that are homozygous in P1 and heterozygous in P2 parents (**Table S4**). These DNA markers were used for construction of linkage map using pseudo-testcross approach (Grattapaglia and Sederoff, 1994). We obtained two linkage maps, one for DNA markers heterozygous in P1 parent, and the other for markers heterozygous in P2 parent **(Fig. S4)**. Since each RAD marker has ∼75 bp sequence, this information was used to associate RAD marker to contigs generated by the de novo assembly (**Fig. S5**). This method allowed us to anchor contigs amounting 378.8 Mb (85.4% of the total genome size) to the linkage maps (**Table S5**) and to combine the two linkage groups and generate pseudochromosomes 1-10 with sizes ranging from 31.5 Mb (Pseudochromosome 5) to 54.6 Mb (Pseudochromosome 1) (**Fig. 2**).

**Fig. 2.**
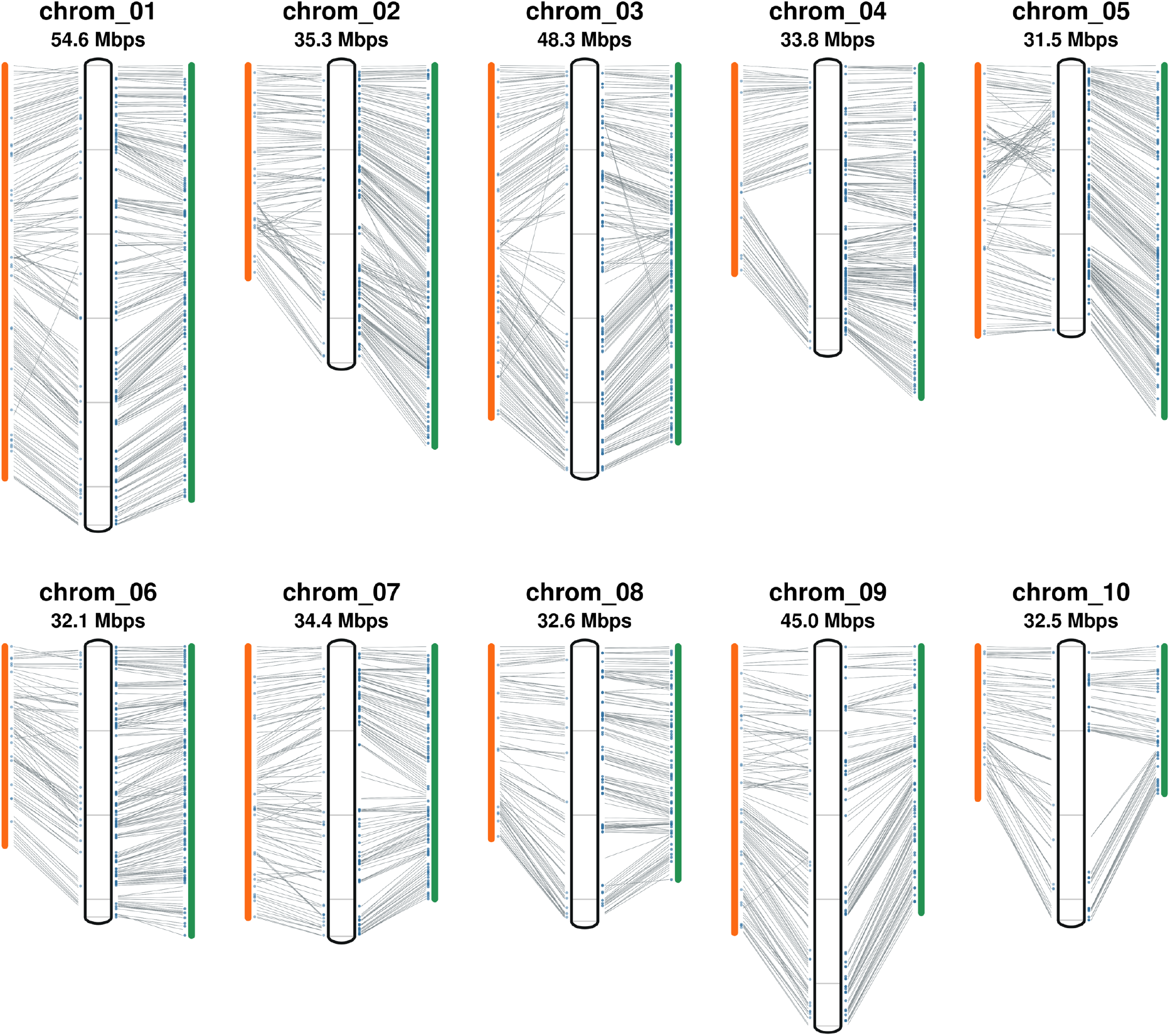
An integrated linkage and physical map of *D. tokoro*. Approximately 85.4% of the *D. tokoro* contig sequences were anchored using a RAD-based genetic map generated with 186 F1 individuals obtained from a cross between Waka1 (P1: female) and Kita1 (P2: male). The 10 pseudochromosomes are numbered from chrom_01 to chrom_10. Markers are located according to genetic distance (cM). The black frame in the center of each group represents the reconstructed pseudochromosome and orange and green bars indicate P1-map and P2-map, respectively. Thin grey lines connecting linkage map and pseudochromosome indicate the positions of markers. The blue dots indicate the positions of PA markers.

BUSCO analysis (Mosè et al. 2020) showed that complete BUSCO value of 98%, indicating that our *D. tokoro* genome sequence is of a sufficient quality as the reference (**Table 1**).

### Gene prediction

We performed RNA-seq of 18 samples representing different organs and developmental stages of *D. tokoro* (**Table S6**). The total size of RNA-seq reads amounted 31.17 Gb. These RNA-seq reads were mapped to the contigs, revealing a total of 29,084 genes, among which 25,447 genes were assigned to pseudochromosomes (**Table 2**).

**Table 2.** Summary of predicted genes in the *D. tokoro* genome

|  | Contigs<br>(2,931) | Pseudochrom.<br>(01-10) |
| --- | --- | --- |
| No. genes | 29,084 | 25,447 |
| (total transcript variant) | (54,847) | (48,271) |
| ORF status |  |  |
| Complete | 21,610 | 19,069 |
| 5' partial | 195 | 166 |
| 3' partial | 4,752 | 4,091 |
| Internal | 157 | 127 |
| No ORF | 2,370 | 1,994 |
| Prediction software |  |  |
| TACO | 24,148 | 21,239 |
| Spaln2 | 1,900 | 1,581 |
| StringTie | 3,036 | 2,627 |

### Sex determination in *D. tokoro*

The 186 F1 progeny derived from a cross between Waka1 female (P1) and Kita1 male (P2) parents segregated in 38 female, 89 male, and 59 non-flowering in 2011 (**Table S3**). We attempted to identify genomic region that shows association with sex of the F1 individuals. As a result of Fisher’s exact test based on the sex and genotype contingency table of each progeny, we found a significant association of the middle position of Pseudochromosome 3 with sex when the DNA markers heterozygous in the male parent (P2) were used. By contrast, there was no association detected if we use the markers heterozygous in the female parent (P1; **Fig. 3**). This result indicates a male heterogametic sex determination (XY) system in *D. tokoro*, and supports our previous analysis with AFLP markers (Terauchi and Kahl, 1999).

**Fig. 3.**
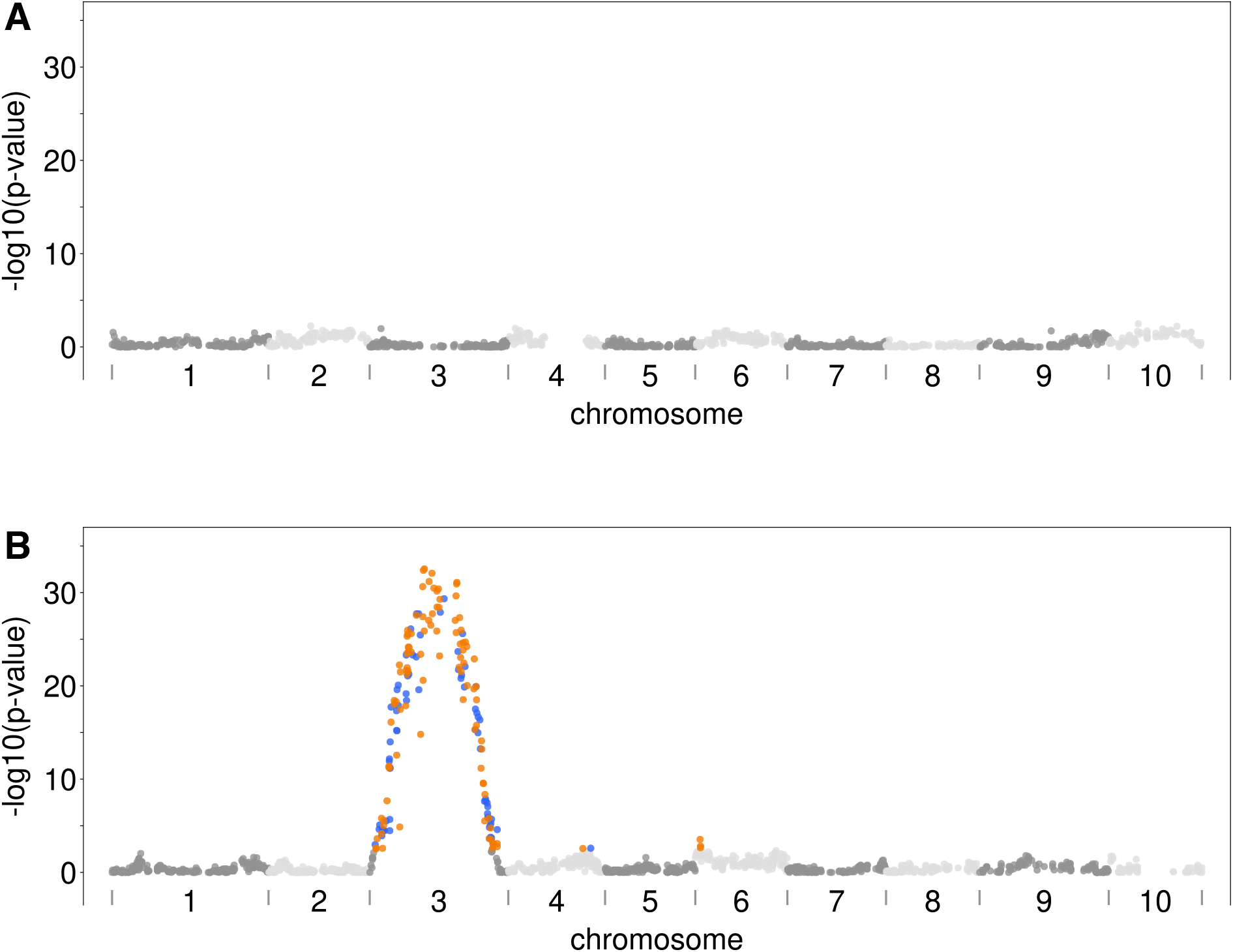
Genome-wide association mapping of sex in the F1 progeny derived from a cross between Waka1 (P1: female) and Kita1 (P2: male) in *D. tokoro*. Manhattan plot of markers associated with sex phenotype as determined by Fisher’s exact test with (A) P1-heterozygous marker set and (B) with P2-heterozygous marker set. Orange and blue dots indicate SNP and presence/absence markers, respectively, showing significant association with sex based on a 5 % false discovery rate (q < 0.05).

## Discussion

Here we report *D. tokoro* draft genome sequence of 443.5 Mb in size. For the species of the genus *Dioscorea*, whole genome sequences are available for *D. rotundata* (Tamiru, 2017; Sugihara et al. 2020), *D. dumetorum* (Siadjeu et al. 2020) and *D. alata* (Bredeson et al. 2022). Genome sizes of these species were 570 Mb (*D. rotundata*), 485 Mb (*D. dumetorum*) and 480 Mb (*D. alata*). The genome of *D. tokoro* is slightly smaller than the genomes of *Dioscorea* species so far reported.

Basic chromosome number of *Dioscorea* is suggested to be ten. *D. tokoro* revealed to have ten linkage groups in this study, which is in line with the report of linkage group obtained by AFLP analysis (Terauchi and Kahl, 1999). It is contrasting to *D. rotundata* (2n = 2x = 40) and *D. alata* (2n = 2x = 40), both belonging to the section Enantiophyllum. It is likely that during the evolution of *Dioscorea*, chromosome duplication occurred. However, no signature of genome duplication observed in *D. rotundata* genome (Tamiru et al. 2017), suggesting that genome duplication happened in a remote past.

Sex determination of *D. tokoro* was confirmed to be XY type male heterogametic system. It is similar to the XY system in *D. alata* (Cormier et al. 2019), but in contrast to ZW female heterogametic system in *D. rotundata* (Tamiru et al. 2017). It is likely that sex determination locus has shifted multiple times in the genus. Future study will identify the genes involved in *D. tokoro* sex determination.

In summary, we determined a draft genome sequence of a wild yam species *D. tokoro*. This chromosome level sequence information will serve as a platform to understand population genomics of this obligate outcrossing species and to elucidate the mechanism and evolution of its sex determination system.

## Materials and Methods

### Plant Materials

A female plant, Waka1 (original code: DT49) (**Fig. S2A**; **Fig. S2D**), was collected from Tahara, Wakayama Pref. in Central Japan. A male plant, Kita1 (original code: 110628-5), was collected from Waga-Sennin of Iwate Pref. in Northern Japan (**Fig. S2B**; **Fig. S2D**). To construct a linkage map, we obtained F1 seeds derived from a cross between Waka1 (P1) and Kita1 (P2) in 2011. We started growing 206 F1 individuals in 2012 and obtained sex data for 186 F1 individuals in 2014 and 2015 (**Fig. S2C**).

### Flow cytometry

Flow cytometry (FCM) analysis was carried out using nuclei prepared from fresh leaf samples. Nuclei were isolated and stained with propidium iodide (PI) and analyzed using a Cell Lab QuantaTM SC Flow Cytometer (Beckman Coulter, USA) following the manufacturer’s protocol.

### Whole genome sequencing and *de novo* assembly

To generate *Dioscorea tokoro* reference genome sequence, we sequenced the male plant Kita1 using the PromethION sequencer (Oxford Nanopore Technologies). First, Kita1 DNA was extracted from fresh leaves as described in our previous report (Tamiru et al., 2017). The extracted DNA was subjected to size selection and purification with a gel extraction kit (Large Fragment DNA Recovery Kit; Zymo Research). Finally, the purified DNA was sequenced by PromethION at GeneBay company, Yokohama, Japan (http://genebay.co.jp). As the first step for genome assembly, we removed the lambda phage genome from Nanopore fastq using NanoLyse 1.1.0 (De Coster et al. 2018) and filtered out reads with an average read quality score less than seven and those shorter than 1,000 bases with Nanofilt v2.2 (De Coster et al. 2018). We also performed two types of Illumina sequencing, 251 bp paired-end sequencing using MiSeq and 150 bp paired-end sequencing using HiSeq4000. Next, we assembled the filtered long DNA sequence reads with the hybrid assembler MaSuRCA v3.3.4 (Zimin et al. 2013), run internally by Flye assembler v2.6 (Kolmogorov et al. 2019).

### Assessing genome completeness

To evaluate the completeness of the gene set in the assembled genome, we applied BUSCO analysis (Bench-Marking Universal Single Copy) v5.1.2 (Mosè et al. 2020). We used the default gene search method ‘metaeuk gene search’ instead of the traditional gene search method using AUGUSTUS (Hoff and Stanke, 2013) and TBLASTN (Camacho et al. 2009). We set “genome” as the assessment mode and used embryophyte_odb10 as the lineage datasets.

### Gene prediction and annotation

For gene prediction, we used RNA-seq data from 18 samples of *D. tokoro*, representing seven organs of Kita1 individual (leaves, stems, root apex, rhizome bud, rhizome root, rhizome stem, and rhizome storage) and 11 different flowering stages in female and male *D. tokoro* plants from the wild (**Table S6**). First, according to the manufacturer’s instructions, total RNAs were used to construct cDNA libraries using a TruSeq RNA Sample Prep Kit V2 (Illumina, USA). Then, the bulked cDNA library was sequenced using the Illumina NextSeq500 platform for 75 bp single-end reads. In the fastq quality control step, we first remove adapters, poly(A), and the reads shorter than 50 bp using FaQCs (Lo and Chain, 2014). Subsequently, we removed low-quality bases from the read end (window size = 5, base quality average = 20) and low-quality reads with an average read quality below 20 using PRINSEQ lite 0.20.4 (Schmieder and Edwards, 2011). Quality trimmed reads were aligned to the assembled genome with HISAT2 v2.1 (Kim et al. 2019) with the options “--max-intronlen 15000 --dta”. Next, transcript alignments were assembled with StringTie v1.3.6 (Pertea et al. 2015) separately for each BAM file. Finally, these GFF files were integrated with TACO v0.7.3 (Niknafs et al. 2017) with the option “--filter-min-length 90”, generating 24,148 gene models within the assembled genome (**Table 2**). Additionally, 34,539 peptide sequences that were predicted in *D. rotundata* genome (Tamiru et al., 2017) [ENSEMBL (http://plants.ensembl.org/Dioscorea_rotundata/Info/Index).] were aligned to assembled genome with Spaln2 v2.3.3 (Iwata and Gotoh, 2012). Consequently, 1,900 CDSs that did not overlap with the new gene models were added to the new gene models (**Table 2**). In addition, the 3,036 transcripts that were assembled in the StringTie program but rejected in the TACO program were added manually. Finally, gene models shorter than 75 bases were removed, and InterProScan v5.36 (N) was used to predict ORFs (open reading frames) and strand information for each gene model. We predicted 29,084 genes, including 54,847 transcript variants (**Table 2**). For gene annotation, the predicted gene models were searched in the Pfam protein family database using InterProScan (Blum et al. 2021) and with the blastx command in BLAST+ (Camacho et al. 2009) with the option “-evalue 1e-10”, using the Viridiplantae database from UniProt as the target database. The resulting gene models and annotations were uploaded to https://genome-e.ibrc.or.jp/resource/dioscorea-tokoro/.

### Identification of parental line-specific heterozygous markers

#### RAD sequencing

We performed RAD-seq to develop the linkage map as previously described (Tamiru et al., 2017). Genomic DNA was extracted from fresh leaves of Waka1, Kita1, and 186 F1 individuals and digested with the restriction enzymes PacI and NlaIII to prepare libraries used to generate 75-bp paired-end reads by Illumina NextSeq500. We remove adapters and the unpaired reads using FaQCs and PRINSEQ lite as previously described. The filtered RAD-seq reads were used as RAD-tags (**Fig. S3**).

#### SNP-type heterozygous markers

RAD-tags were aligned to the assembled genome of *D. tokoro* in this study using BWA (ver. 0.7.12). SNP-based genotypes for P1, P2, and F1 individuals was obtained as a variant call format (VCF) file. The VCF file was generated from BAM files of P1, P2, and F1 individuals using SAMtools (ver 1.5), and the VCF variants were called and filtered using BCFtools (ver 1.5). As a result, 5,894 P1-or P2-heterozygous SNP markers were selected (shown as “All RAD markers” in **Table S4**). Next, to increase the accuracy of the selected markers, their segregation (1:1 ratio) was confirmed in F1 individuals obtained from a cross between P1 and P2. If the segregation ratio was out of the confidence interval (*P* < 0.001) hypothesized by the binomial distribution, *B* (n = number of individuals, *P* = 0.5), the markers were excluded from further analysis. Finally, 3,057 P1-heterozygous SNP markers and 1,559 P2-heterozygous SNP markers were selected (shown as “Confirmed segregation ratio” in **Table S4**). Additional details are provided in the **Supplementary Method**.

#### Presence/absence-type heterozygous markers

The presence/absence-type markers were defined based on the alignment depth of parental line RAD-tags. The presence/absence-type markers were called by the following method: First, the VCF file was generated from BAM files of P1 and P2 and selected the region where either P1 or P2 had sufficient read depth (≥ 8) and that the other parental line had no read depth in that region. Next, BEDtools (ver 2.26) converted continuous positions in the VCF file to a feature, and only sufficiently wide features (width ≥ 50 bp) were retained as the BED file. For these regions in the BED file, the F1 individual’s genotypes were classified into three categories (depth ≥ 3, depth = 0, others) and three genotypes (“presence,” “absence,” “NA”). As a result, 5,071 PA markers were selected (shown as “All RAD markers” in **Table S4**). We then applied the same binomial test as for the SNP-type heterozygous markers. Finally, 480 P1-heterozygous PA markers and 1,682 P2-heterozygous PA markers were selected (shown as “Confirmed segregation ratio” in **Table S4**). Additional details are provided in the **Supplementary Method**.

#### Integration of SNP-type and presence/absence-type heterozygous markers

We integrated SNP-type and PA-type heterozygous markers to develop parental line-specific linkage maps. Two types of markers were defined: P1-heterozygous markers and P2-heterozygous markers. If an SNP-type marker was heterozygous in P1 but homozygous in P2 or if a PA-type marker was present in P1 and absent in P2, it was classified as a P1-heterozygous marker set. Conversely, if a SNP-type marker was homozygous and heterozygous in P1 and P2, respectively, or if a PA-type marker was absent in P1 but present in P2, it was classified as a P2-heterozygous marker set.

### Linkage maps construction

#### Pruning and flanking markers by Spearman’s correlation coefficients

Pairwise matrix of Spearman’s correlation coefficients (ρ) were calculated for every maker pair in each contig in each marker set (P1-heterozygous marker set and P2-heterozygous marker set). According to the histogram of absolute ρ calculated from each contig, most markers on the same contigs were correlated with each other. Therefore, we pruned correlated flanking markers to remove redundant markers. Finally, we obtained 2,818 markers for linkage mapping (shown as “Pruning and flanking” in **Table S4**).

#### Linkage mapping

We converted the flanking markers obtained as described in the previous section into the genotype-formatted data for constructing genetic linkage maps using MSTmap (Wu, 2008) with following parameter sets: “populationtype DH; distancefunction kosambi; cutoffpvalue 0.000000000001; nomapdist 15.0; nomapsize 0; missingthreshold 25.0; estimationbeforeclustering no; detectbaddata no; objective_function ML” for P1-heterozygous marker set and P2-heterozygous marker set. After trimming the orphan linkage groups, we solved the complemented-phased duplex linkage groups caused by coupling-type and repulsion-type markers in the pseudo-testcross method. Finally, two parental-specific linkage maps were constructed. These two linkage maps were designated as P1-map (constructed using P1-heterozygous marker set) and P2-map (constructed using P2-heterozygous marker set) (**Fig. S4A; Fig. S4C**). The order and names of each linkage group were organized according to the P2-map (**Fig. S4 and Fig. S5**). The linkage groups were visualized by R/qtl (Broman et al., 2003).

### Generation of pseudochromosomes

Based on a matrix derived from the contigs shared between the P1- and P2-maps, i.e., linkage groups (**Fig. S5**), the contigs were anchored and linearly ordered as pseudochromosomes. First, we identified contigs whose markers were allocated to different linkage groups during the anchoring and ordering process. Such contigs were further divided into sub-contigs to ensure that they were not allocated to wrong pseudochromosomes. Next, we divided the contigs at the proper positions as described previously (Tamiru et al. 2017). Finally, we followed the described method (Tamiru et al. 2017) to generate ten pseudochromosomes.

### Identification of sex associated region

To identify the sex-associated genomic region, we performed Fisher’s exact test using the genotype of 127 F1 individuals based on 2,730 markers located on the pseudochromosomes (**Table S4**) and their sex phenotype (**Table S3**). Fisher’s exact test was performed using the fisherexact function in the python SciPy package. A significance threshold of 5% false discovery rate (FDR) was calculated using the multipletests function in the python statsmodels package with the option method= “fdr_bh” (Benjamini / Hochberg procedure).

## Data Availability

All sequencing read data generated for this work have been deposited at the DNA Databank of Japan (DDBJ) database under BioProject PRJDB12945; see **Table S1** and **S2** for individual sample accession numbers. The genomic sequence file (fasta), gene annotation file (gff3), and gene/protein sequences file (fasta) are available at the following URL: https://genome-e.ibrc.or.jp/resource/dioscorea-tokoro/

## Acknowledgments

We dedicate this work to Günter Kahl, a pioneer of *Dioscorea* molecular genetics. In this research work we used the NIG supercomputer at ROIS National Institute of Genetics, and the supercomputer of ACCMS, Kyoto University.

---

**Fig. S1.**
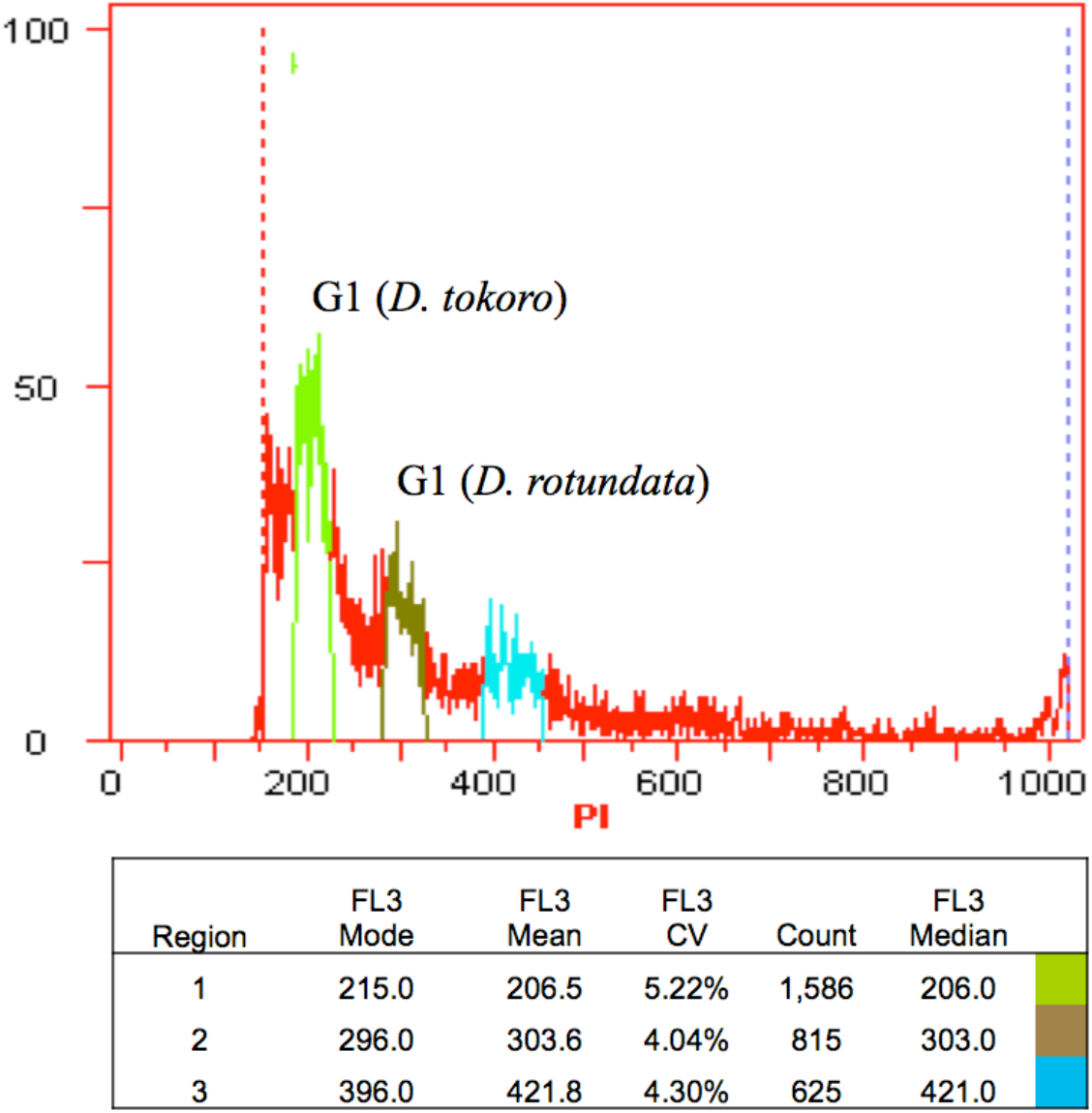
Size estimation of *D. tokoro* genome by flow cytometry. Flow cytometry analysis was carried out using nuclei prepared from fresh leaf samples of a wild plant of *D. tokoro* collected in Kitakami, Iwate, Japan and a plant of *D. rotundata* maintained in a greenhouse at Iwate Biotechnology Research Center (IBRC). *D. rotundata* (570 Mb) was served as an internal reference standard of known genome size (Tamiru et al. 2017). Nuclei were isolated and stained with propidium iodide (PI) and analyzed using a Cell Lab QuantaTM SC Flow Cytometer (Beckman Coulter, USA) following the manufacturer’s protocol. The ratio of G1 peak mean [*D. tokoro* (206.5): *D. rotundata* (303.6) = 0.680] was used to estimate the genome size of *D. tokoro* to be 388 Mb (570 Mb × 0.68).

**Table S1.** Summary of filtered ONT reads.

|  |  |
| --- | --- |
| Number of reads<br>(before filtering) | 2,515,235<br>(3,126,676) |
| Total base-pairs (Gbp)<br>(before filtering) | 27.4<br>(32.7) |
| Genome coverage* (x) | 70.6 |
| Mean read length (kbp) | 10.9 |
| Longest fragment (kbp) | 11.0 |
| Shortest fragment (bp) | 1 |
| Accession No. | DRR344532 |

\*Genome coverage is estimated from the expected genome size of *D. tokoro* (388 Mb).

**Table S2.** Summary of non-filtered Illumina short reads.

| Sequence platform | Read length<br>(bp) | Total base<br>pairs(Gbp) | Genome coverage | Accession No. |
| --- | --- | --- | --- | --- |
| Illumina MiSeq | 35-251 | 24.6 | 63.4x | DRR344531 |
| Illumina HiSeq 4000 | 150 | 37.8 | 97.4x | DRR347075 |
|  | Total | 62.4 | 160.8x |  |

**Table S3.** Number of male, female and non-flowering progeny derived from a Waka1 (P1) x Kita1 (P2) cross.

|  |  | Female | Male | Not<br>flowered |
| --- | --- | --- | --- | --- |
| Parent | Waka1 (P1) | 1 |  |  |
|  | Kita1 (P2) |  | 1 |  |
| Progeny | 186<br>individuals | 38 | 89 | 59 |

**Fig. S2.**
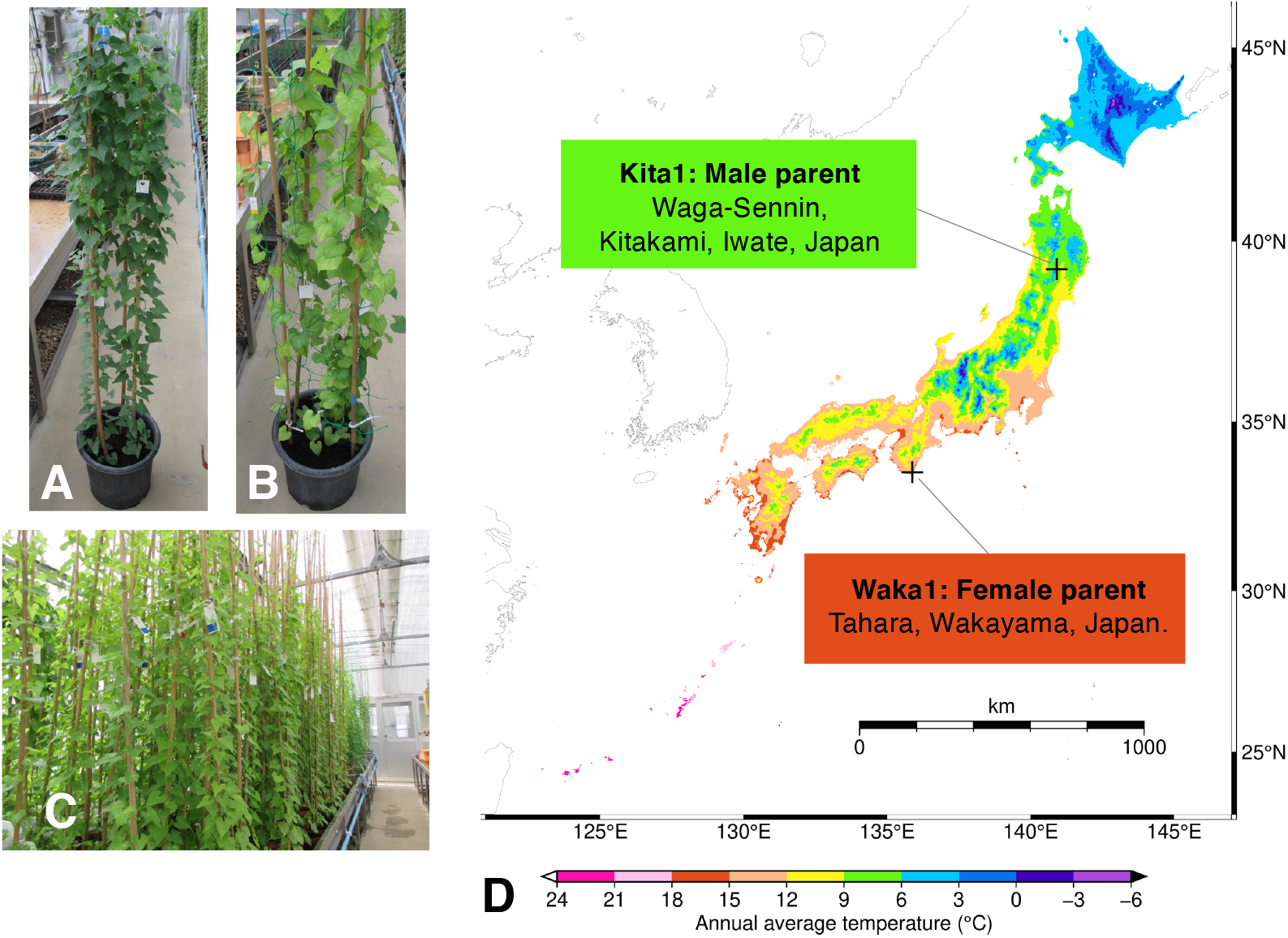
*D. tokoro* plants used for a genetic cross and information of sites of origin. (A) A female individual Waka1. (B) A male individual Kita1. (C) Sideview of 186 F1 individuals obtained from a cross between Waka1 and Kita1. (D) Waka1 was collected from Tahara, Wakayama Pref. in the Kinki district of Japan. This place is close to the coast. The latitude and longitude are 33.538, 135.860, respectively, and the average annual temperature is 15-18°C. Kita1 was collected from Waga-Sennin in Kitakami, Iwate Pref. in the Tohoku district of Japan. This place is a mountainous. The latitude and longitude are 39.295, 140.896, respectively, and the average annual temperature is 6-9 °C. This map was created with GMT, Version 6.1.1 (Wessel et al. 2019). The annual average temperature on the map of Japan is drawn using “Annual average (climate) mesh data”, downloaded from the National Land Numerical Information Download Service (JPGIS2.1) (https://nlftp.mlit.go.jp/ksj/index.html) published by the Ministry of Land, Infrastructure, Transport and Tourism.

**Fig S3.**
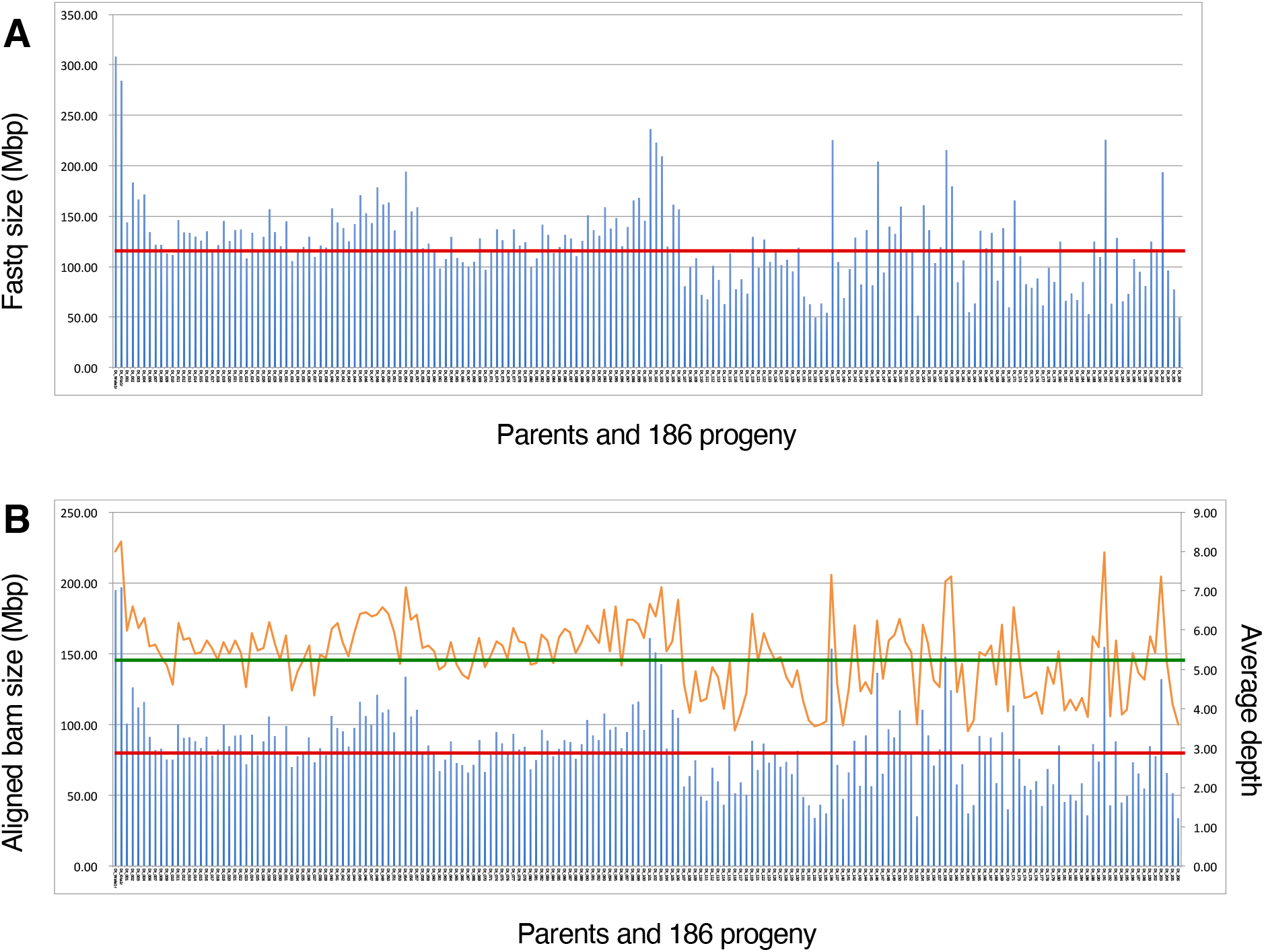
Summary of RAD tags generated for 186 F1 individuals derived from a cross between Waka1 (P1: female) and Kita1 (P2: male). In all graphs, the two bars on the left end indicate the parents Waka1 and Kita1, and the other indicate 186 F1 individuals. (A) The total size of filtered fastq of each individual (blue bars). The horizontal red line indicates the average fastq size of the progeny (120.7 Mbp). (B) Aligned bam size (blue bars) and average read depth at genomic regions in the reference genome aligned by the RAD-tags (orange line). The horizontal red line indicates the average aligned bam size of the progeny (82.3 Mbp), and the horizontal green line indicates the average read depth of the progeny (5.34 Mbp).

**Table S4.** Number of RAD markers used for anchoring the contigs after filtering.

| **Filtering step** | **Type** | **LG** | **SNP** | **PA** | **total** |
| --- | --- | --- | --- | --- | --- |
| All RAD markers | testcross |  | **5,894** | **5,071** | **10,965** |
| Confirmed segregation ratio | P1-hetero |  | 3,057 | 480 | 3,537 |
|  | P2-hetero |  | 1,559 | 1,682 | 3,241 |
|  | **(total)** |  | **4,616** | **2,162** | **6,778** |
| Pruning and flanking | P1-hetero | 49 | 988 | 181 | 1,169 |
|  | P2-hetero | 55 | 768 | 881 | 1,649 |
|  | **(total)** |  | **1,756** | **1,062** | **2,818** |
| Anchoring contigs | P1-hetero | 10 | 946 | 180 | 1,126 |
|  | P2-hetero | 10 | 724 | 880 | 1,604 |
|  | **(total)** |  | **1,670** | **1,060** | **2,730** |

**Fig. S4.**
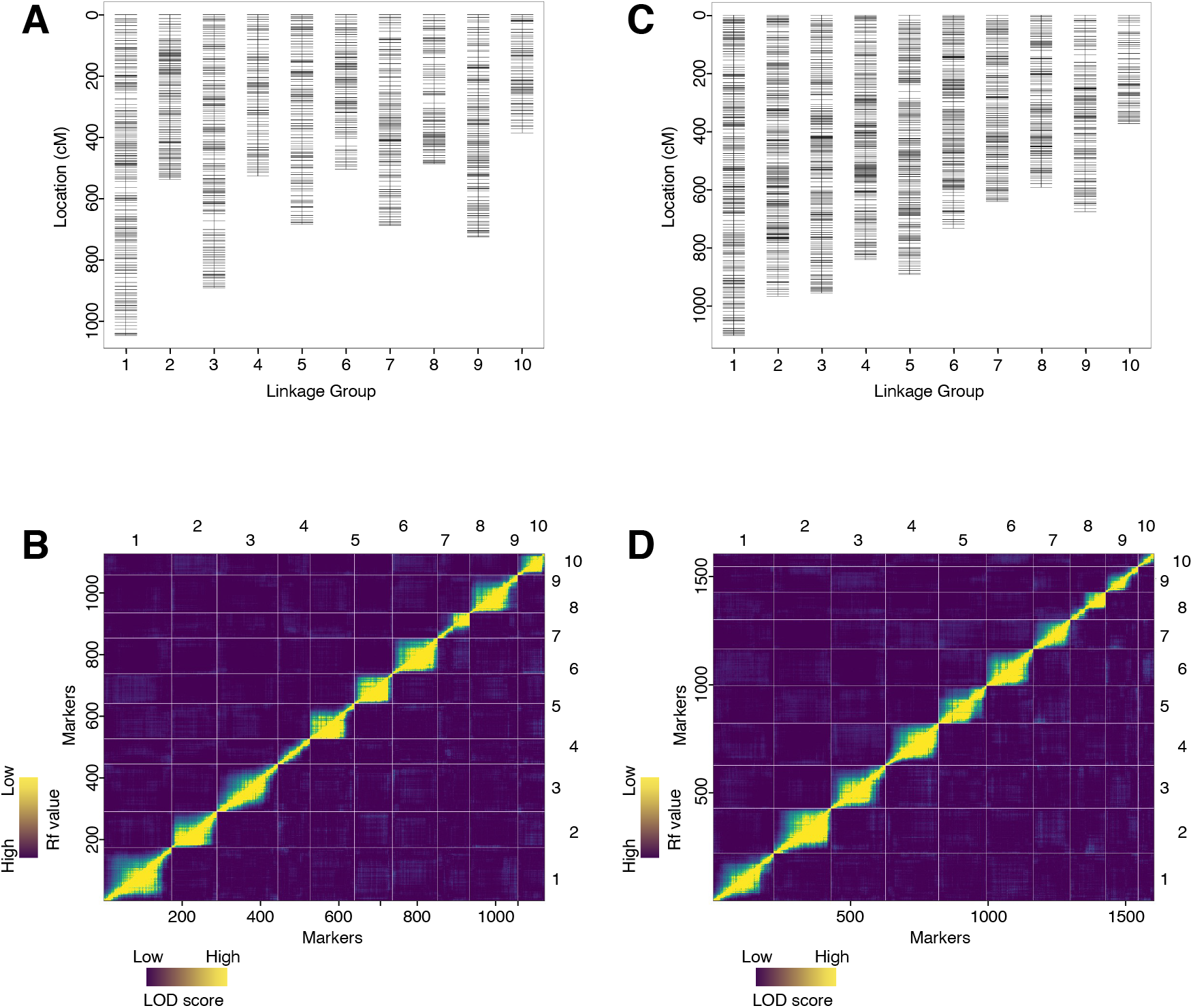
RAD-seq-based linkage map of *D. tokoro* generated by the pseudo-testcross method using 186 F1 individuals. (A) P1-map generated using P1-heterozygous marker set. (B) Plots of estimated recombination fractions (upper-left triangle) and LOD score (lower-right triangle) for P1-Map. (C) P2-map generated using P2-heterozygous marker set. (D) Plots of estimated recombination fractions (upper-left triangle) and LOD score (lower-right triangle) for P2-Map. Yellow indicates linked (large LOD score or small recombination fraction) and blue indicates not linked (small LOD score or large recombination fraction).

**Fig. S5.**
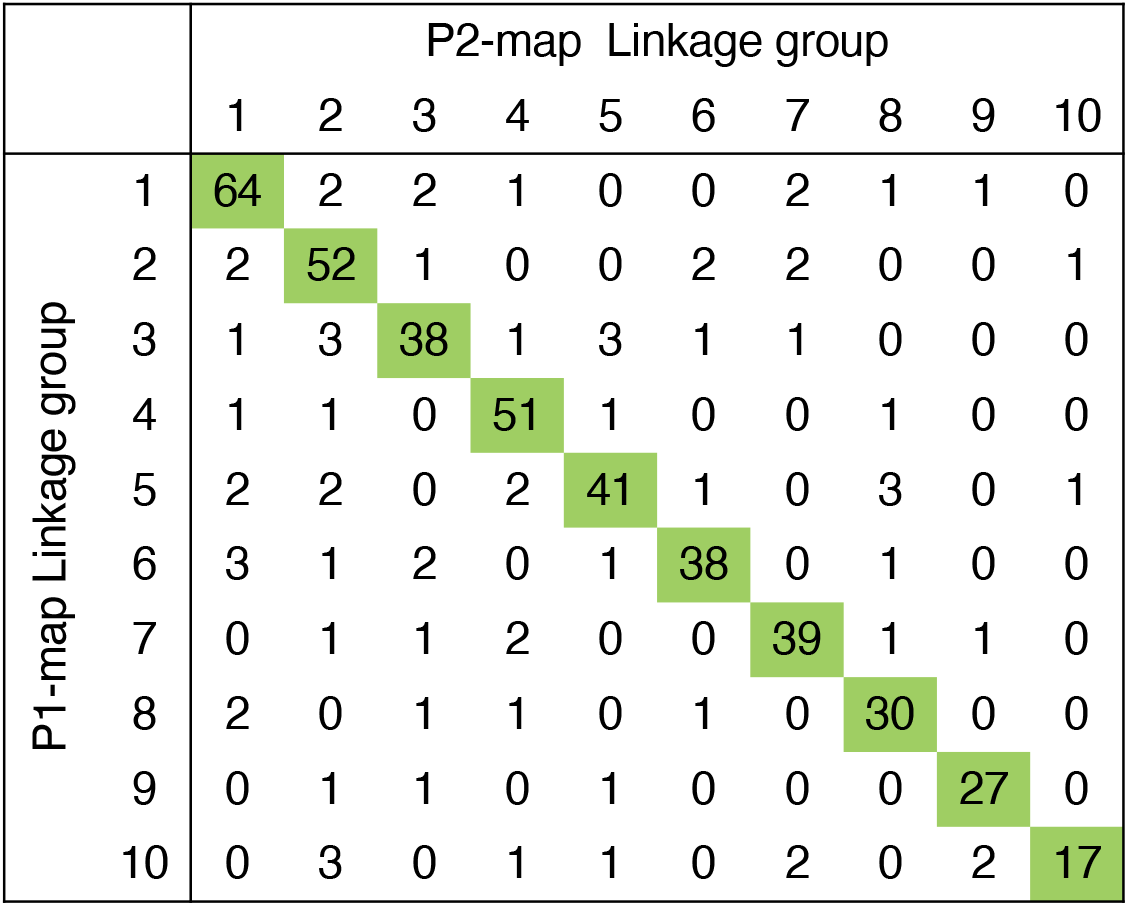
A matrix of the number of shared contigs between the P1-map and P2-map. Both P1-Map and P2-Map contained 10 LGs.

**Table S5.** Summary of contigs anchored by RAD markers of the 10 linkage groups.

|  | P1<br>hetero | P2<br>hetero | P1 P2<br>total |
| --- | --- | --- | --- |
| Markers in 10 LGs | 1,126 | 1,604 | 2,730 |
| contig information<br>anchored by marker: |  |  |  |
| number | 566 | 963 | 1,123 |
| contig number % | 19.3 | 32.9 | 38.3 |
| contig total bps | 303,270,609 | 320,556,813 | 378,798,395 |
| contig total % | 68.4 | 72.3 | 85.4 |

\*The reference fasta size is 443,501,147 bp.

**Table S6.** Details of 18 RNA-seq samples.

| No. | Organ | Phase | Sex | Stage* | Collected material | Fastq size |
| --- | --- | --- | --- | --- | --- | --- |
| 1 | leaves |  | male |  | Kita1 | 1.71 |
| 2 | stems |  | male |  | Kita1 | 1.35 |
| 3 | root apex |  | male |  | Kita1 | 1.66 |
| 4 | rhizome bud |  | male |  | Kita1 | 1.86 |
| 5 | rhizome root |  | male |  | Kita1 | 1.61 |
| 6 | rhizome stem |  | male |  | Kita1 | 1.79 |
| 7 | rhizome storage |  | male |  | Kita1 | 1.78 |
| 8 | shoot apex | vegetative | unknown |  | multiple wild | 1.62 |
| 9 | shoot apex | reproductive | female | 0 | multiple wild | 1.87 |
| 11 | shoot apex | reproductive | female | 1 | multiple wild | 1.70 |
| 13 | shoot apex | reproductive | female | 2 | multiple wild | 1.81 |
| 10 | shoot apex | reproductive | male | 0 | multiple wild | 1.71 |
| 12 | shoot apex | reproductive | male | 1 | multiple wild | 1.74 |
| 14 | shoot apex | reproductive | male | 2 | multiple wild | 1.91 |
| 15 | mature bud |  | female |  | multiple wild | 1.85 |
| 16 | mature bud |  | male |  | multiple wild | 1.69 |
| 17 | flower |  | female |  | multiple wild | 1.83 |
| 18 | flower |  | male |  | multiple wild | 1.83 |
|  |  |  |  |  | total | 31.17 |

\*The reproductive shoot stage are indicated as follows. 0: shoot apical meristem (SAM), 1: inflorescence with unopened bracts, 2: inflorescences below 10 mm with unseparated bottom buds.

## Supplementary Method

### Identification of parental line-specific heterozygous RAD markers

#### Heterozygous SNP markers

SNP genotypes for P1, P2, and F1 progenies were obtained as a VCF file. The VCF file was generated as follows: (i) SAMtools v1.5 mpileup command with the option “-t DP,AD,SP -B -Q 18 -C 50”; (ii) BCFtools v1.5 call command with the option “-P 0 -v -m -f GQ,GP”; (iii) BCFtools view command with the options “-i ‘INFO/MQ≥40, INFO/MQ0F≤0.1, and AVG(GQ)≥10”; and (iv) BCFtools norm command with the option “-m+any.” We rejected the variants with low read depth (< 10) or low genotype quality scores (< 10) in the two parents. Likewise, we regarded variants with low read depth (< 8) or low genotype quality scores (< 5) in F1 progenies as missing and only retained the variants with low missing rates (< 0.3). Subsequently, only bi-allelic SNPs were selected by the BCFtools view command with the option “-m 2 -M 2 -v snps”. Referring to the genotypes in the VCF file, heterozygous genotypes called by unbalanced allele frequency (out of 0.1-0.9 in F1 progenies) were regarded as missing, and filtering for missing rate (< 0.1) was applied again. As a result, 5,894 P1-or P2-heterozygous SNP markers were selected (shown as “All RAD markers” in Table S5). Next, a binomial test was performed to reject SNPs affected by segregating distortion in the F1 progenies. This binomial test assumes that the probability of success rate is 0.5 based on the two-side hypothesis, and we regarded variants having p-value less than 0.001 as segregation distortion. Finally, 3,057 P1-heterozygous SNP markers and 1,559 P2-heterozygous SNP markers were selected (shown as “Confirmed segregation ratio” in Table S5).

#### Heterozygous presence/absence RAD markers

The presence/absence markers were defined based on the alignment depth of parental line RAD-tags. A VCF file was generated to search for positions with contrasting read depth between the two parental plants P1 and P2 using the following commands: (i) SAMtools mpileup command with the option “-B -Q 18 -C 50”; (ii) BCFtools call command with the option “-A”; and (iii) BCFtools view command with the options “-i ‘MAX(FMT/DP)≥8 and MIN(FMT/DP)≤0’ -g miss -V indels”. This means that one of the parents (P1 or P2) has enough read depth (≥ 8) and another parent has no reads aligned on that region. Subsequently, we converted continuous positions in the VCF file to a feature that provides a region’s start and end coordinate information using the BEDTools v.2.26 merge command with the option “-d 10 -c 1 -o count”. We only retained sufficiently wide features (≥ 50 bp) in the BED file. Using the depth value in each feature given in the BED file, presence/absence (PA) -based genotypes for parental plants P1 and P2 and F1 progenies were determined. For P1 and P2, we regarded genotypes having depth ≥ 4 as present genotypes, meaning the heterozygosity of presence and absence, while those having depth = 0 were classified as absent genotypes, meaning the homozygosity of absence. For F1 progenies, we classified markers with depth > 2 and = 0 as present and absent markers, respectively. Referring to the genotypes, heterozygous genotypes called by unbalanced allele frequency (out of 0.1-0.9 in F1 progenies) were rejected. As a result, 5,071 PA markers were selected (shown as “All RAD markers” in Table S5). Next, we applied the same binomial test for PA heterozygous markers as SNP-type heterozygous markers. Finally, 480 P1-heterozygous PA markers and 1,682 P2-heterozygous PA markers were selected (shown as “Confirmed segregation ratio” in Table S5).

